# In silico comparative genomics of SARS-CoV-2 to determine the source and diversity of the pathogen in Bangladesh

**DOI:** 10.1101/2020.07.20.212563

**Authors:** Tushar Ahmed Shishir, Iftekhar Bin Naser, Shah M. Faruque

## Abstract

The COVID19 pandemic caused by SARS-CoV-2 virus has severely affected most countries of the world including Bangladesh. We conducted comparative analysis of publicly available whole-genome sequences of 64 SARS-CoV-2 isolates in Bangladesh and 371 isolates from another 27 countries to predict possible transmission routes of COVID19 to Bangladesh and genomic variations among the viruses. Phylogenetic analysis indicated that the pathogen was imported in Bangladesh from multiple countries. The viruses found in the southern district of Chattogram were closely related to strains from Saudi Arabia whereas those in Dhaka were similar to that of United Kingdom and France. The 64 SARS-CoV-2 sequences from Bangladesh belonged to three clusters. Compared to the ancestral SARS-CoV-2 sequence reported from China, the isolates in Bangladesh had a total of 180 mutations in the coding region of the genome, and 110 of these were missense. Among these, 99 missense mutations (90%) were predicted to destabilize protein structures. Remarkably, a mutation that leads to an I300F change in the nsp2 protein and a mutation leading to D614G change in the spike protein were prevalent in SARS-CoV-2 genomic sequences, and might have influenced the epidemiological properties of the virus in Bangladesh.

## Introduction

The pandemic of coronavirus disease referred to as COVID-19 pandemic, which originated in Wuhan, China in December 2019 is ongoing and has now spread to 213 countries and territories. As of July 2020, the pandemic has caused about 16 million cases and over half a million death. This novel virus of the Coronaviridae family and Betacoronavirus genus (1, 2) designated as severe acute respiratory syndrome coronavirus 2 (SARS-CoV-2), is the causative agent of COVID-19. Previously two other coronaviruses, namely SARS-CoV and MERS-CoV have demonstrated high pathogenicity and caused epidemic with mortality rate ∼10% and ∼34% respectively affecting more than 25 countries each time (3–5). However, SARS-CoV-2 has proven to be highly infectious and caused pandemic spread to over 213 countries and territories. Besides its devastating impact in North America and Europe, the disease is now rapidly spreading in South America including Brazil, and in South Asian countries particularly India, Pakistan and Bangladesh (6).

The virus was first detected in Bangladesh in March 2020 (7). Although infections remained low until the end of March it began to rise steeply in April. By the end of June, new cases in Bangladesh grew to nearly 150,000 and the rate of detection of cases compared to the total number of samples tested increased to about 22% which was highest in Asia (6).

SARS-CoV-2 is a positive-sense single-stranded RNA virus with a genome size of nearly 30kb. The 5’ end of the genome codes for a polyprotein which is further cleaved to viral non-structural proteins whereas the 3’ end encodes for structural proteins of the virus including surface glycoproteins spike (S), membrane protein (M), envelop protein (E) and nucleocapsid protein (N) (8). Like other RNA viruses, SARS-CoV-2 is also inherently prone to mutations due to high recombination frequency resulting in genomic diversity (9–11). Due to the rapid evolution of the virus, development of vaccines and therapies may be challenging. To monitor the emergence of diversity, it is important to conduct comparative genomics of viruses isolated over time and in various geographical locations. Comparative analysis of genome sequences of various isolates of SARS-CoV-2 would allow to identify and characterize the variable and conserved regions of the genome and this knowledge may be useful for developing effective vaccines, as well as in molecular epidemiological tracking. Thousands of SARS-CoV-2 virus genomes have been sequenced and submitted in public databases for further study. This include 66 SARS-CoV-2 genomic sequences submitted from Bangladesh in the Global initiative on sharing all influenza data (GISAID) database, till 11th June 2020 (12). We conducted comparative analysis of publicly available genome sequences of SARS-CoV-2 from 27 countries to predict the origin of viruses in Bangladesh by studying a time-resolved phylogenetic relationship. Later, we analyzed the variants present in different isolates of Bangladesh to understand the pattern of mutations in relation to the ancestral Wuhan strain, find unique mutations, and possible effect of these mutations on the stability of encoded proteins, and selection pressure on genes.

## Materials and methods

A total of 435 whole genome sequences of SARS-CoV-2 including 64 sequences isolated in Bangladesh (detail information in S1 Table), and that of 5 isolates of each month between January to May, 2020 isolated in 27 different countries with high frequency of infection were included in this analysis. Source and number of sequence are presented in Table 1 (detail information are provided in S2 Table). However, since only number of sequences were reported from different African countries, we included all sequences from the African countries and categorized collectively as African sequence (12). Reference sequence included in various analysis was the sequence of the ancestral strain from Wuhan, China (NC_045512.2) (13).

**Table 1.** Number of SARS-CoV-2 genomic sequences reported from different countries included in the study.

| **Country** | **Number of Sequences** |
| --- | --- |
| African region | 17 |
| Australia | 27 |
| Bangladesh | 65 |
| Brazil | 15 |
| Canada | 19 |
| China | 31 |
| France | 20 |
| Germany | 20 |
| India | 18 |
| Iran | 5 |
| Italy | 30 |
| Mexico | 8 |
| Nepal | 1 |
| Pakistan | 3 |
| Russia | 14 |
| Saudi Arabia | 19 |
| Spain | 15 |
| Sri Lanka | 6 |
| Sweden | 5 |
| Turkey | 10 |
| United Kingdom | 31 |
| USA | 55 |

Selected sequences were annotated by Viral Annotation Pipeline and iDentification (VAPiD) tool (14), and multiple sequence alignment was carried out using Mafft algorithm (15). Maximum likelihood phylogenetic tree was constructed with IQ-TREE (16), the generated tree was reconstructed based on time-calibration by TreeTime (17), and visualized on iTOL server (18). For analysis of mutations, sequence were mapped with minimap2 (19), and variants were detected using SAMtools (20) and snp-sites (21). A haplotype network was generated based on mutations in genome using PopArt (22). Sequences were then put into different clades based on specific mutations proposed in GISAID (23) and further classified as D614G type (24, 25). Subsequently, another phylogenetic tree and haplotype network containing only SARS-CoV-2 sequences from Bangladeshi was constructed and categorized using the same tools, and additionally one step further clustered with TreeCluster (26). The direction of selection in sequences from Bangladesh was calculated by the SLAC algorithm (27) in the Datamonkey server (28). Finally, the effects of the mutations on protein stability were predicted using DeepDDG (29).

## Results and Discussion

### Phylogenetic analysis of all SARS-CoV-2 sequences

A total of 435 Genomic Sequences of SARS-CoV-2 reported from various countries (Table 1) which included 64 sequences from Bangladesh and the sequence of the ancestral SARS-CoV-2 isolated in Wuhan, China were analyzed in the time-resolved phylogenetic tree. Sequences from Bangladesh belonged to three different clusters of which one cluster carried 43 of the 64 sequences, and shared the same node with sequence from Germany while they had a common ancestry with isolates from the United Kingdom (Fig 1). The remaining two clusters of SARS-CoV-2 sequences contained 4 and 5 sequences respectively from Bangladesh, and they shared the same node with sequence of SARS-CoV-2 reported from India, and also shared a common ancestry with isolates from Saudi Arabia. Besides, 12 lone sequences that did not belong to any of these clusters were found to have similarity with sequences from Europe including United Kingdom, Germany, France, Italy, and Russia. One of these sequences was closely related to sequence reported from the USA. Subsequently, all SARS-CoV-2 sequences from representative countries were clustered based on some specific mutations sustained, into 7 different clades as mentioned by GISAID. In this analysis, the sequences from Bangladesh were found to be distributed in all clades except V (Figs 1 and 2).

**Fig 1.**
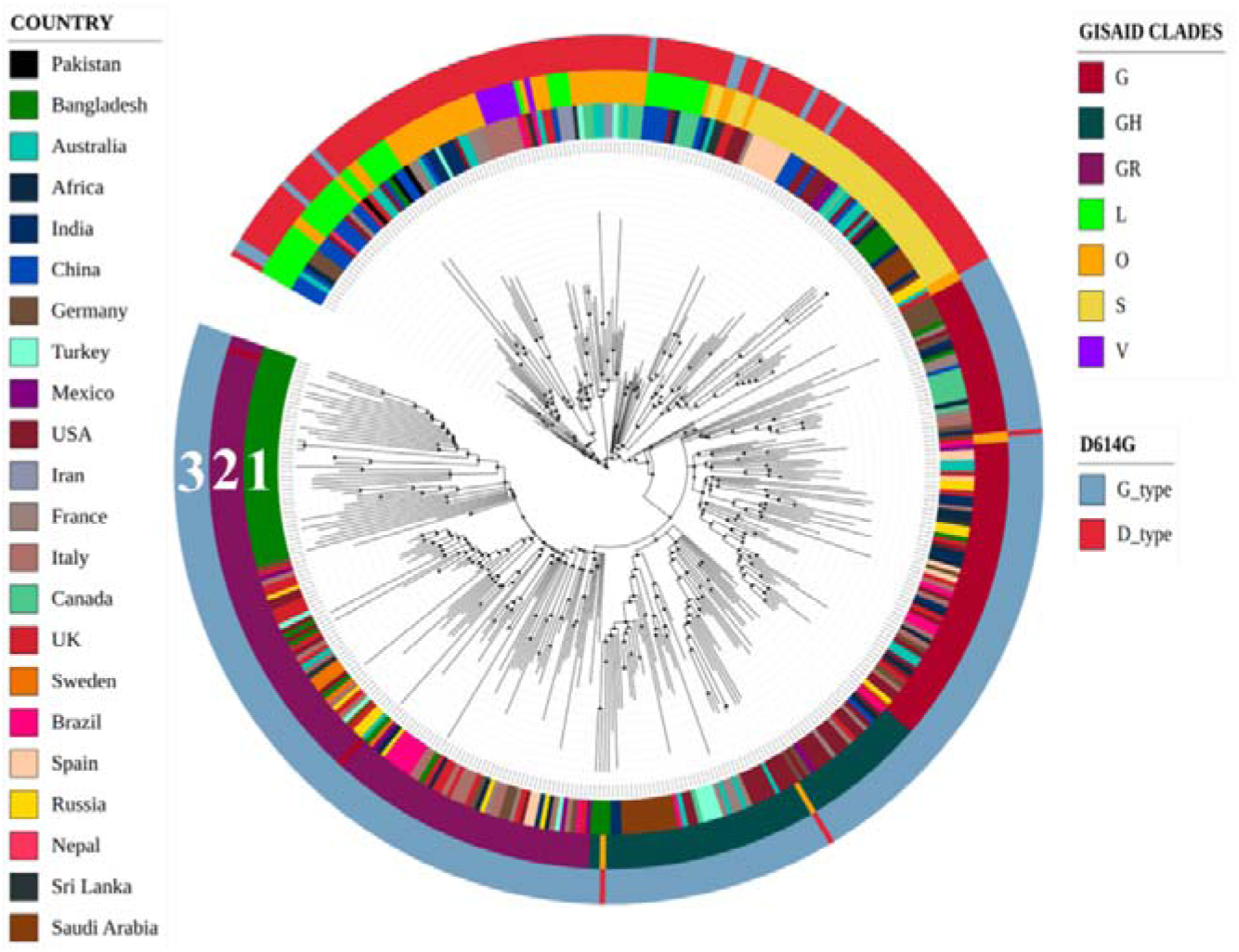
Time-resolved phylogenetic tree of sequences from different countries. Circle 1 exhibits the location of the sequences and how these are distributed; circle 2 demonstrates GISAID clades based on some specific mutations and circle 3 represents D614G classification.

**Fig 2.**
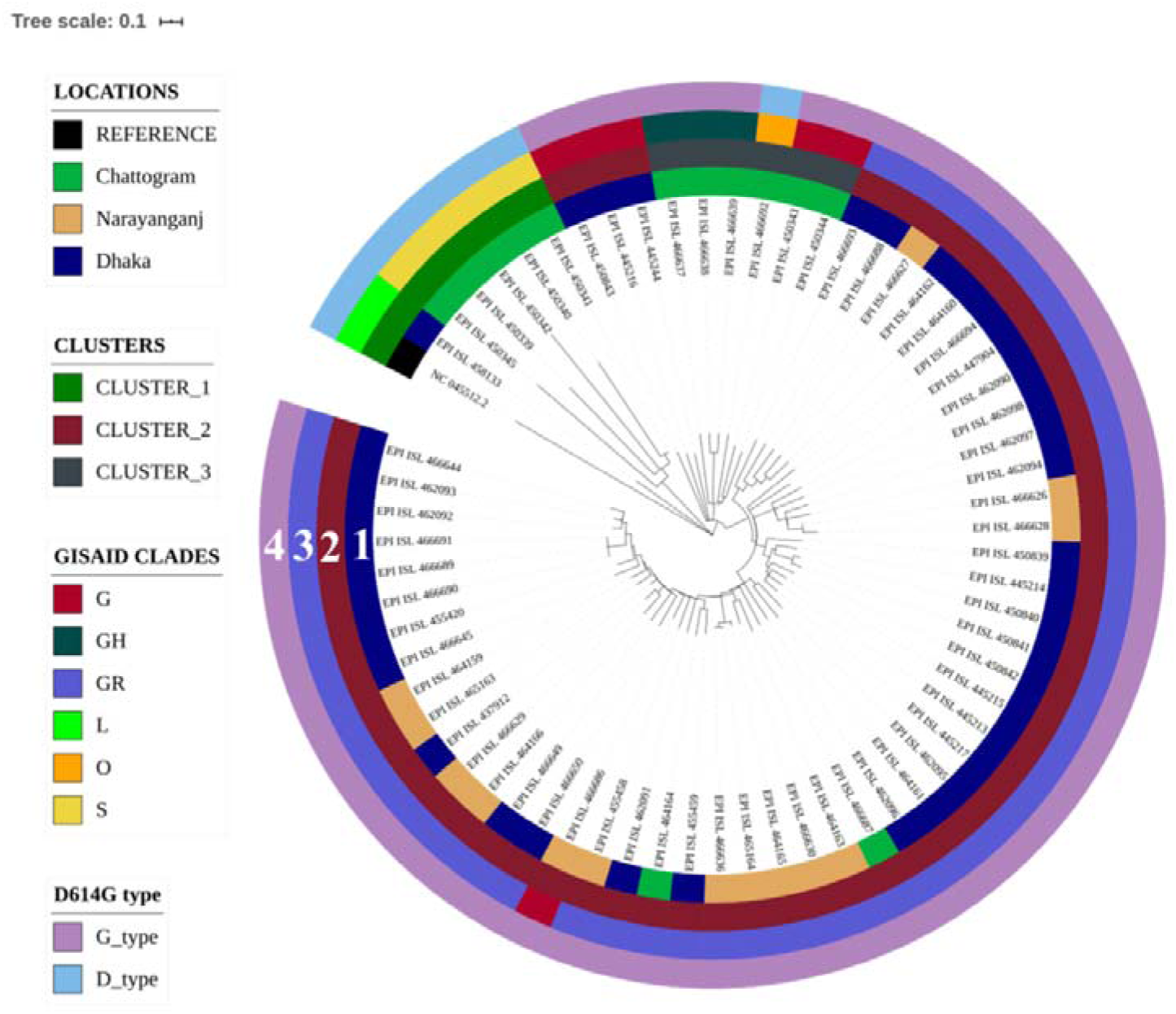
Time-resolved phylogenetic tree of SARS-CoV-2 sequences from Bangladesh. Circle 1 shows the location of isolation of the samples, circle 2 shows the three clusters in which various sequences belong based on their similarity, circle 3 shows GISAID clades of different sequences, and circle 4 represents whether the sequences are D type or G type.

### Phylogenetic comparison of SARS-CoV-2 sequences from Bangladesh

In order to understand the evolutionary relationship and possible transmission dynamics of SARS-CoV-2 in Bangladesh at a higher resolution, another time-resolved phylogenetic tree carrying only sequences of the pathogen isolated in various regions of Bangladesh was generated using the sequence of the first SARS-CoV-2 reported from Yuhan, China as a reference. Of the three clusters produced in this analysis, cluster-1 included mostly isolates from Chattogram and one isolate from Dhaka, cluster-2 included isolates from Dhaka, Narayanganj and Chattogram districts, whereas cluster-3 included isolates from Chattogram only. As mentioned above, the isolates from Bangladesh were found to be distributed in all 7 GISAID clades based on specific mutations, except in clade V (Fig 2). Most isolates of Dhaka and Narayanganj (47 of 52) belonged to the GR clade, whereas those of Chattogram belonged to five different GSID clades (G, GH, GR, O, and S).

The major international airport in Bangladesh is situated in the capital city Dhaka, whereas the major seaport is located in Chattogram. Based on the phylogenetic analysis, all isolates of Dhaka were the descendant of SARS-CoV-2 found in European countries, more specifically France and the United Kingdom. On the other hand, most isolates of Chattogram were found closely related to Saudi Arabian isolates. Moreover, considering the GSID clades, the presence of S clade was absent among Dhaka whereas most isolates of Chattogram was found to belong to the S clade. Clearly these two genomic variants of SARS-CoV-2 were initially imported by travelers from different countries, and the two variants initially spread in the two areas. That the isolates of Narayanganj and two isolates of Dhaka are closely related, indicates that the SARS-CoV-2 strain imported initially through international traveler to Dhaka later spread to Narayanganj, which is a densely populated city with river ports and large business centres.

### D614G mutations in spike protein

The SARS-CoV-2 sequences were also categorized according to D614G type mutation (Fig 1). This particular subtype with a non-silent (Aspartate to Glycine) mutation at 614th position of the Spike protein is presumed to have rapidly outcompeted other preexisting subtypes, including the ancestral strain. The D614G mutation generates an additional serine protease (Elastase) cleavage site near the S1-S2 junction of the Spike protein (25). All but one sequence from Dhaka and Narayanganj were found to be of G type which carries Glycine at position 614 whereas sequences of Chattogram carry sequences of both types (Fig 2). In addition, the first sequence from Bangladesh carried G614 type of surface glycoprotein, which indicate that this dominant variant was present since the first isolation of SARS-CoV-2 in Bangladesh and the mutant virus might have been imported to the country from Europe, and the presence of the mutation might have facilitated viral transmission.

### Haplotype analysis of SARS-CoV-2 sequences

Relationships among DNA sequences within a population are often studied by constructing and visualizing a haplotype network. We constructed a haplotype network by the Median Joining algorithm and found that 338 of 434 SARS-CoV-2 sequences from representative countries were alike, therefore formed a large haplo group (Fig 3A). However, there were presences of a significant number of unique lineages too consisting of a single or multiple SARS-CoV-2 sequences (Fig 3A). This network demonstrated the closeness of the sequences and their pattern of mutation beyond the geographical boundary. Several SARS-CoV-2 isolates appeared to have sustained certain common mutations along with certain unique mutations. Although a large proportion of sequences from Bangladesh belonged to the common cluster (Fig 3A), there was a significant number of unique nodes as well due to mutations overtime subsequent to being carried into Bangladesh (Fig 3A). Therefore analysis of the sequences from Bangladesh provided further insight of their mutation patterns. The haplotype network revealed that viruses isolated in Bangladesh had certain unique mutations in them, and as a result they belonged to different haplo groups and no significant cig group (Fig. 3A). Most of the isolates sustained a significant number of mutations compared with each other. In addition it further confirms that most isolates from Chattogram (Fig 3B) were not directly related to those isolated in Dhaka or Narayanganj.

**Fig 3.**
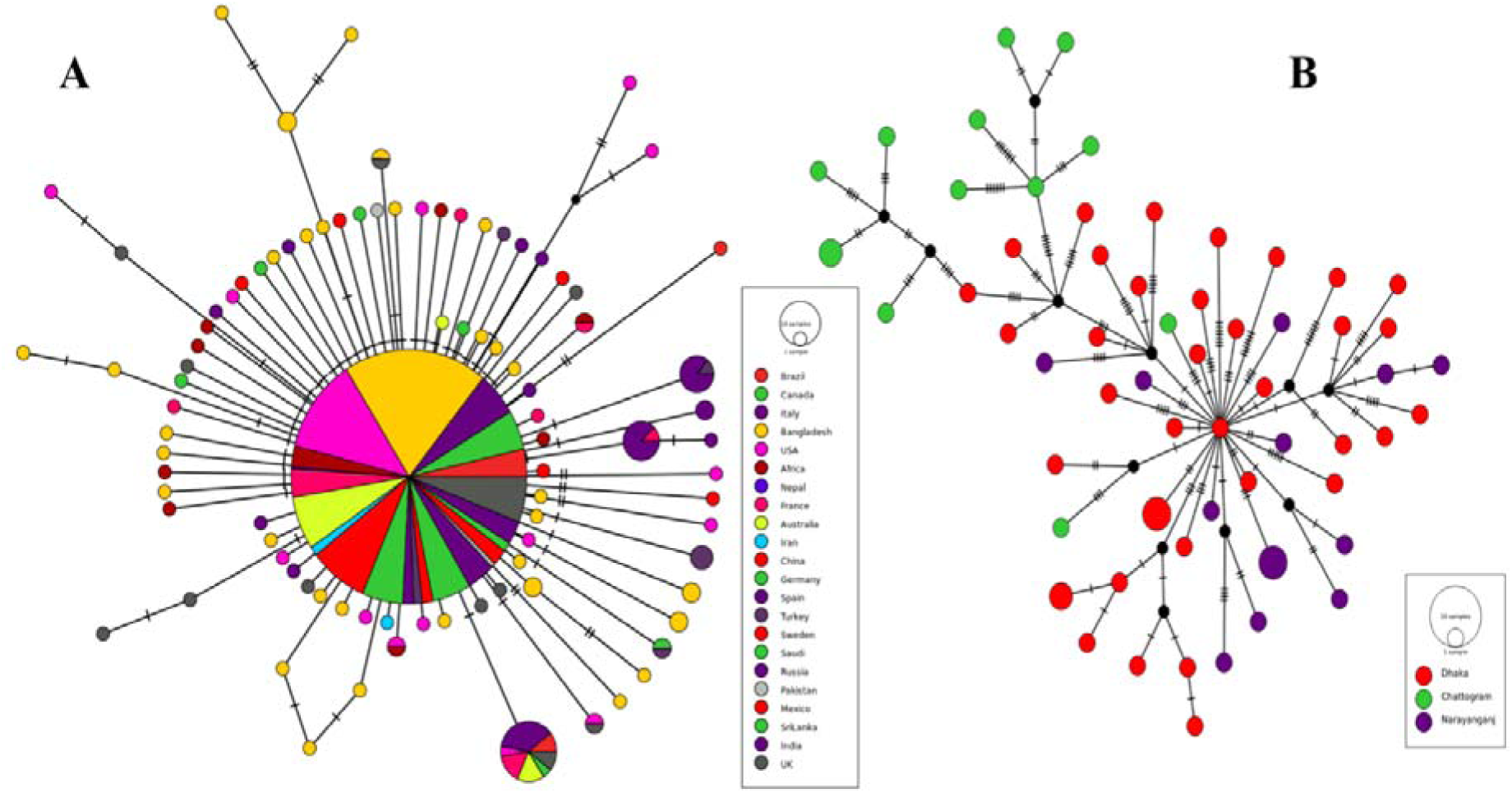
Haplotype network of selected SARS-CoV-2 genome sequences. (A) The largest circle represents a group of sequences from different countries that are similar and few sequences are connected with other sequences through undetermined intermediate due to harboring unique mutations denoted by small black circle. (B) Haplotype network of sequences from Bangladesh; the number of dashes in the connecting line denotes the number of mutations against each other.

### Location and predicted effects of the mutations

We detected the presence of 209 point mutations in 64 SARS-CoV-2 isolates from Bangladesh when compared to the reference sequence from Yuhan, China. In addition, 19 isolates were found to have lost significant portions of their genome, and as a result lost sequences for some non-structural proteins such as ORF7 and ORF8 while other deletions were upstream or downstream gene variants (S3 Table). Among the point mutations, 29 mutations were in the non-coding region of the genome and 180 were in coding regions. Ten of the 29 non-coding mutations were in upstream non-coding region and rest was in downstream non-coding region of the genome. Seventy mutations in the coding region were synonymous and 110 mutations predicted substituted amino acids. Among twelve predicted ORFs, ORF1ab which comprises approximately 67% of the genome encoding 16 nonstructural proteins had more than 60 percent of the total mutations while gene E encoding envelope protein and ORF7b were conserved and did not carry any mutation. Though ORF1 harbored the highest number of mutations, mutation density was highest in ORF10 considering ORF lengths. Details and distribution of the mutations are presented in Table 2 and full analysis report is placed in S3 table.

**Table 2.**
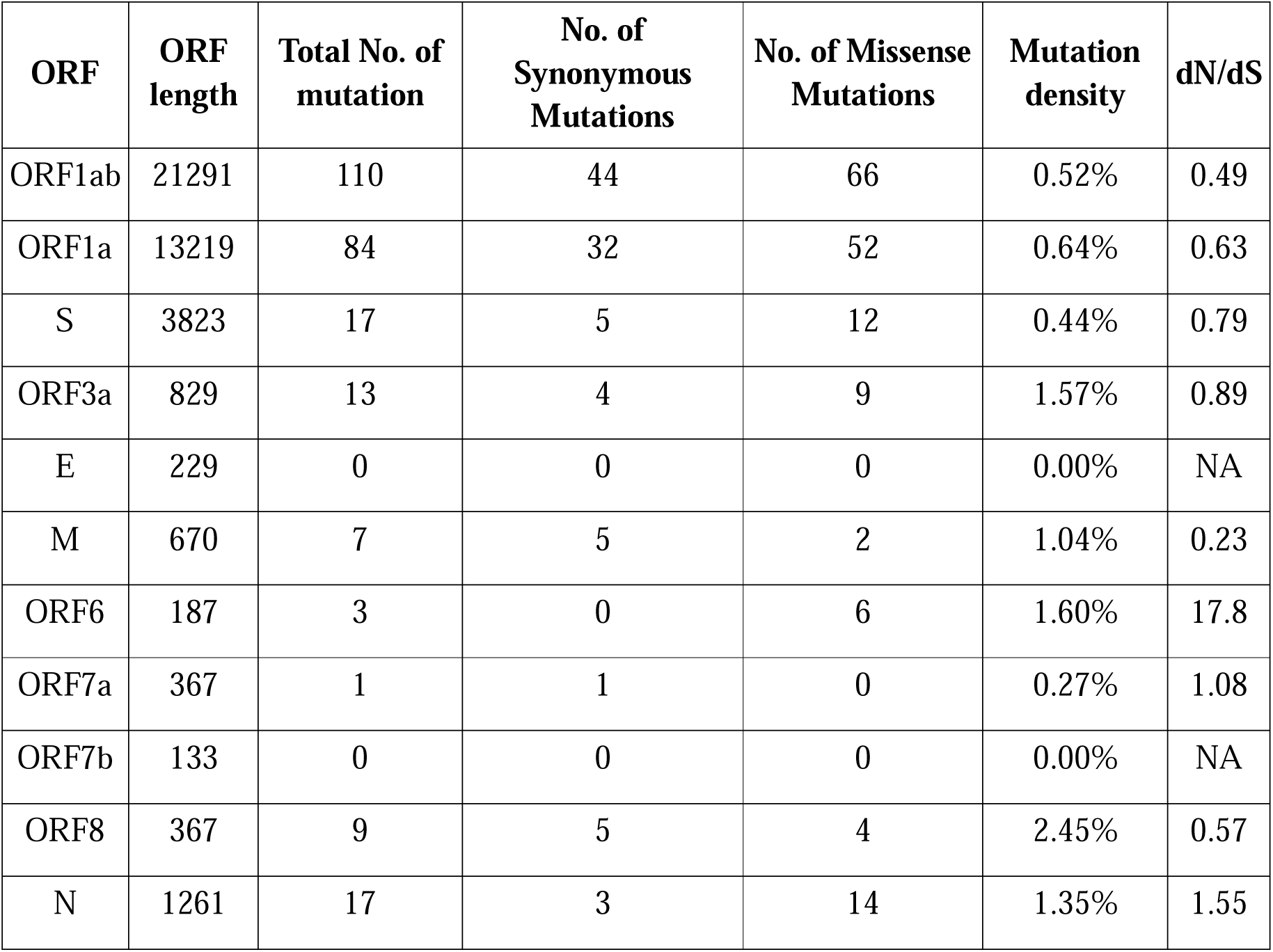
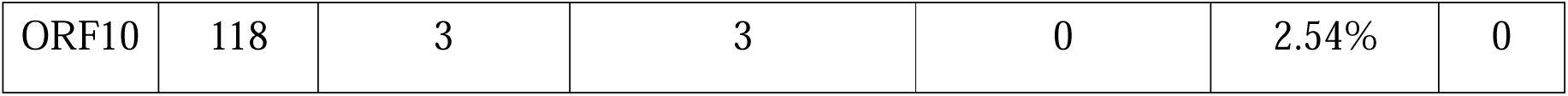
Count of mutation in each gene, types of mutation, mutation density and directional selection value. (dN/dS> 1 is positive selection, dN/dS = 1 neutral selection, dN/dS< 1 negative selection, dN/dS = 0 is conserved region)

| ORF | ORF length | Total No. of mutation | No. of Synonymous Mutations | No. of Missense Mutations | Mutation density | dN/dS |
| --- | --- | --- | --- | --- | --- | --- |
| ORF1ab | 21291 | 110 | 44 | 66 | 0.52% | 0.49 |
| ORF1a | 13219 | 84 | 32 | 52 | 0.64% | 0.63 |
| S | 3823 | 17 | 5 | 12 | 0.44% | 0.79 |
| ORF3a | 829 | 13 | 4 | 9 | 1.57% | 0.89 |
| E | 229 | 0 | 0 | 0 | 0.00% | NA |
| M | 670 | 7 | 5 | 2 | 1.04% | 0.23 |
| ORF6 | 187 | 3 | 0 | 6 | 1.60% | 17.8 |
| ORF7a | 367 | 1 | 1 | 0 | 0.27% | 1.08 |
| ORF7b | 133 | 0 | 0 | 0 | 0.00% | NA |
| ORF8 | 367 | 9 | 5 | 4 | 2.45% | 0.57 |
| N | 1261 | 17 | 3 | 14 | 1.35% | 1.55 |
| ORF10 | 118 | 3 | 3 | 0 | 2.54% | 0 |

In sequences from Bangladesh, 241C>T and 3037C>T changes were the two most abundant mutations found in 58 out of 64 isolates, and often found simultaneously (Table 3). Position 241 is located in the non-coding region whereas the mutation in position 3037 was synonymous. On the other hand, 57 sequences were found to harbor 14408C>T and 23403A>G mutations which altered amino acid Pro>Leu and Asp>Gly respectively, and these two mutations were found to be present simultaneously as well. In addition, other co-evolving mutations found were (241C>T, 3037C>T, 14408C>T, 23403A>G), (28881G>A, 28882G>A, 28883G>C), (8782C>T, 28144T>C), (4444G>T, 8371G>T, 29403A>G). Apart from highly abundant mutations, several less common and unique mutations were also present in the sequences analyzed (Table 2, Fig 4).

**Table 3.** High-frequency mutations present in SARS-CoV-2 sequences from Bangladesh, and their predicted effect on the stability of protein structures. (ddG< 0: decrease stability, ddG> 0: increase stability)

| Position | Gene | Feature | Amino acid change | Frequency | ddG (kcal/mol) |
| --- | --- | --- | --- | --- | --- |
| 241 C > T | 5' UTR | Non-coding | - | 58/64 |  |
| 1163 A > T | ORF1ab | missense | I300F | 44/64 | -0.756 |
| 3037 C > T | ORF1ab | Synonymous | - | 58/64 |  |
| 14408 C > T | ORF1ab | Missense | P4715L | 57/64 | 1.463 |
| 23403 A > G | S | Missense | D614G | 57/64 | -0.45 |
| 28881 G > A | N | Missense | R203L | 49/64 | -3.4 |
| 28882 G > A | N | Synonymous | - | 49/64 |  |
| 28883 G > C | N | Missense | G204A | 49/64 | -4.3 |

**Fig 4.**
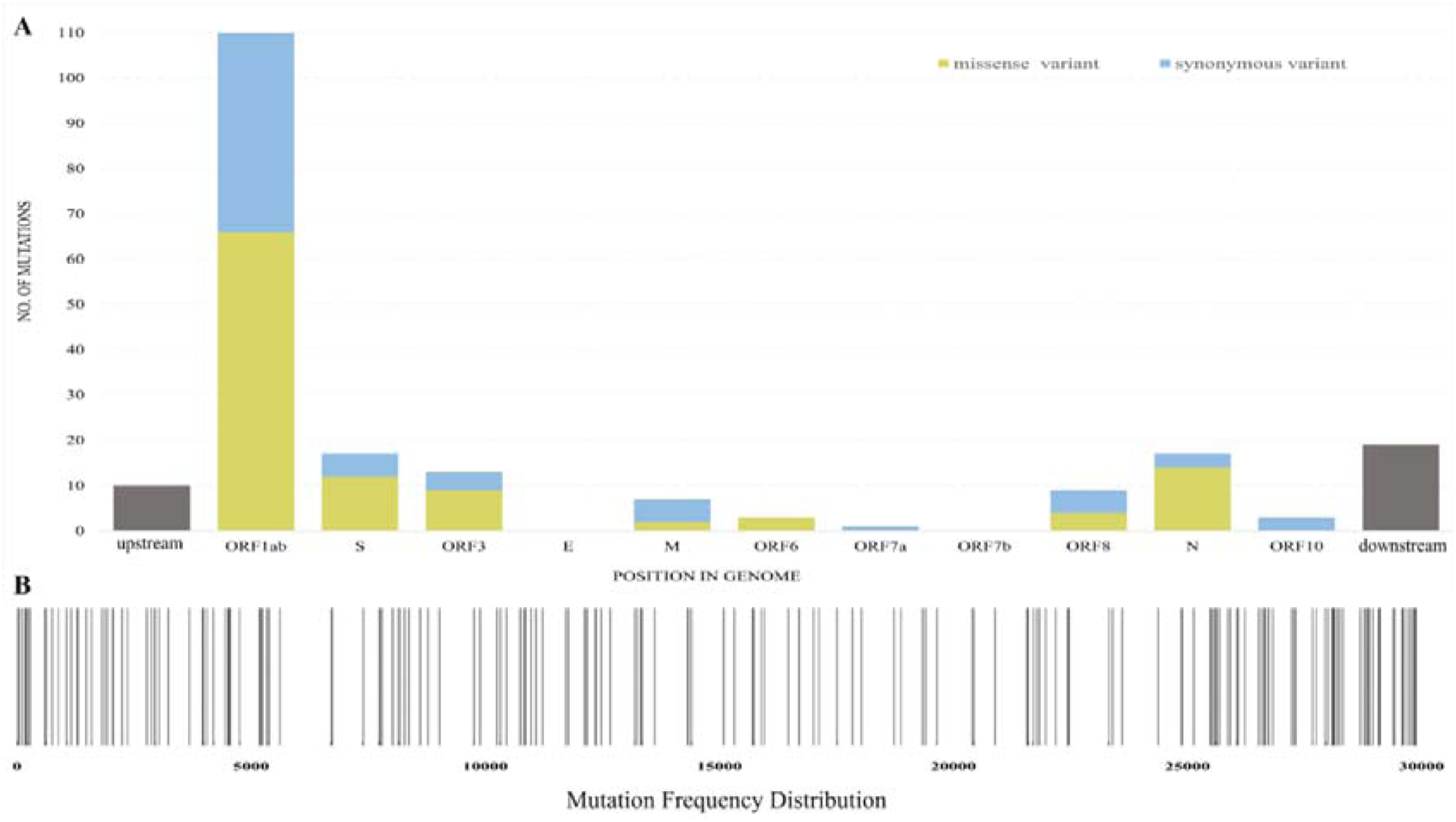
Mutations present in SARS-COV-2 genome sequences reported from Bangladesh. (A) Number of mutations at different regions (B) Mutation distribution frequency

As a result of continuous mutations genes are seemingly going through natural selection. ORF6 was predicted to have dN/dS value of 17.8 due to the presence of higher number of missense than synonymous mutations. This finding indicates that ORF6 is rapidly evolving and is highly divergent. The ORF6 protein is an accessory protein whose function is yet to be fully elucidated (30). The ORF7a and Nucleocapsid phosphoprotein had dN/dS values 1.08 and 1.55 respectively which confer their strong evolution to cope up with challenges under positive selection pressure. ORF10 is predicted to be conversed with dN/dS value 0 while envelope protein and ORF7b did not harbor any mutation and was conserved. On the other hand, ORF3a and surface glycoprotein might approach toward positive selection pressure and evolve but rests of the proteins were under negative selection pressure.

Among 110 missense mutations, 99 were predicted to destabilize the corresponding proteins while only 11 mutations predictably increase stability. Ten protein stabilizing mutations were present in ORF1 with 1 mutation in ORF3a. On the other hand, destabilizing mutations were distributed among most of the ORFs (Table 4). None of the mutations in structural proteins were predicted to increase stability.

**Table 4.** Mutations present in different ORFs present in Bangladeshi sequences.

| **ORF** | **Unique Mutation**<br>**in SARS-Cov-2 in**<br>**Bangladesh** | **Remarks** |
| --- | --- | --- |
| ORF1ab | 93 | A mutation at 1163A>T (Ile300Phe) in ORF1 operon which lower the structural stability of the protein was found existing in 44 strains of Bangladesh but absent in representative countries |
| S | 13 | A highly prevalent mutation 23403A>G (Asp614Gly) which ease the viral transmission, dominating throughout the world was found to be present in 57 of 64 sequences in Bangladesh. |
| ORF3a | 9 | A mutation at 25563G>T (Gln57His) was found highly prevalent among selected sequences from the USA was also present in Bangladeshi sequences. |
| M | 4 | Unique mutations were synonymous |
| ORF6 | 3 | 3 missense mutations were absent in sequences from representative countries. |
| ORF7a | 1 | Synonymous mutation |
| ORF8 | 7 | 3 missense mutations but their frequency was lower |
| N | 10 | Mutations at 28881G>A (Arg203Lys), 28882G>A (Arg203ARG), 28883G>C (Gly204Arg) were harbored in 49 of 64 sequences which were found highly prevalent among viruses in European and North American countries |
| ORF10 | 3 | All mutations were synonymous and found this gene conserved |

In summary, mutation analysis revealed point mutations as well as deletion of base pairs. Deletions of the base pairs were associated with missing non-structural proteins and predictably affected certain viral properties since ORF7a protein is the growth factor of the coronavirus family, induce apoptosis, and promotes viral encapsulation (31–33) while ORF8 is associated with viral adaptation by playing role in host-virus interaction (34, 35). Furthermore, we have found that some genes are under positive selection pressure indicating that the virus is fast-evolving presumably to evade host cell’s innate immunity; which should be taken into special consideration prior to vaccine development or other treatment strategies. Finally, a missense mutation at 1163A>T changing the amino acid isoleucine to phenylalanine in Nsp2 protein was found uniquely among 44 isolates in Bangladesh. Nsp2 is a methyltransferase like domain that interacts with PHB and PHB2, and modulates the host cell survival strategy by affecting cellular differentiation, mitochondrial biogenesis, and cell cycle progression to escape from innate immunity (35, 36). This unique and high-frequency mutation might be a further interest of study, considering death rate against the infection rate in Bangladesh.

## Supporting information

S1_SEQUENCES FROM BANGLADESH

S2_SEQUENCES FROM SELECTED COUNTRIES

S3_MUTATIONS IN BANGLADESHI SEQUENCES

## Supporting information

**S1 Table. Sequences from Bangladesh**.

**S2 Table. Sequences from selected countries.**

**S3 Table. Mutations in Bangladeshi sequences**.

## Notes

### Competing Interest Statement

The authors have declared no competing interest.

## References

1. WHO. Coronavirus disease COVID-2019 - Situation Report 169. World Heal Organ. 2020; Available from https://www.who.int/emergencies/diseases/novel-coronavirus-2019/situation-reports

2. Lefkowitz EJ, Dempsey DM, Hendrickson RC, Orton RJ, Siddell SG, Smith DB. Virus taxonomy: The database of the International Committee on Taxonomy of Viruses (ICTV). Nucleic Acids Res. 2018;46(D1):D708–17.

3. Zhong NS, Zheng BJ, Li YM, Poon LLM, Xie ZH, Chan KH, et al. Epidemiology and cause of severe acute respiratory syndrome (SARS) in Guangdong, People’s Republic of China, in February, 2003. Lancet. 2003;362(9393):1353–1358.

4. De Wit E, Van Doremalen N, Falzarano D, Munster VJ. SARS and MERS: recent insights into emerging coronaviruses. Nature Reviews Microbiology. 2016;14(8):523–534.

5. Yin Y, Wunderink R. MERS, SARS and other coronaviruses as causes of pneumonia. Respirology. 2017;23(2):130–137.

6. Webmeter. Coronavirus Age, Sex, Demographics (COVID-19) - Worldometer. 2020. Available from: www.worldometers.info

7. Rahaman Khan MH, Hossain A. COVID-19 Outbreak Situations in Bangladesh: An Empirical Analysis. medRxiv. 2020. Forthcoming

8. Cui J, Li F, Shi Z. Origin and evolution of pathogenic coronaviruses. Nature Reviews Microbiology. 2019;17(3):181–192.

9. Denison MR, Graham RL, Donaldson EF, Eckerle LD, Baric RS. Coronaviruses: An RNA proofreading machine regulates replication fidelity and diversity. RNA Biology. 2011; 8(2):270–279.

10. Yang Y, Liu C, Du L, Jiang S, Shi Z, Baric RS, et al. Two Mutations Were Critical for Bat-to-Human Transmission of Middle East Respiratory Syndrome Coronavirus. J Virol. 2015; 89(17):9119–9123.

11. Andersen KG, Rambaut A, Lipkin WI, Holmes EC, Garry RF. The proximal origin of SARS-CoV-2. Nature Medicine. 2020; 89(17):9119–9123.

12. Shu Y, McCauley J. GISAID: Global initiative on sharing all influenza data – from vision to reality. Eurosurveillance. 2017;22(13).

13. Wu F, Zhao S, Yu B, Chen YM, Wang W, Song ZG, et al. A new coronavirus associated with human respiratory disease in China. Nature. 2020; 579(7798):265–269.

14. Shean RC, Makhsous N, Stoddard GD, Lin MJ, Greninger AL. VAPiD: A lightweight cross-platform viral annotation pipeline and identification tool to facilitate virus genome submissions to NCBI GenBank. BMC Bioinformatics. 2019;20(1).

15. Katoh K, Standley DM. MAFFT multiple sequence alignment software version 7: Improvements in performance and usability. MolBiolEvol. 2013;30(4):772–780.

16. Nguyen LT, Schmidt HA, Von Haeseler A, Minh BQ. IQ-TREE: A fast and effective stochastic algorithm for estimating maximum-likelihood phylogenies. MolBiolEvol. 2015; 32(1):268–274.

17. Sagulenko P, Puller V, Neher RA. TreeTime: Maximum-likelihood phylodynamic analysis. Virus Evol. 2018;4(1).

18. Letunic I, Bork P. Interactive Tree Of Life (iTOL) v4: recent updates and new developments. Nucleic Acids Res. 2019; 47(W1):W256–W259.

19. Li H. Minimap2: Pairwise alignment for nucleotide sequences. Bioinformatics. 2018; 34(18):3094–3100.

20. Li H, Handsaker B, Wysoker A, Fennell T, Ruan J, Homer N, et al. The Sequence Alignment/Map format and SAMtools. Bioinformatics. 2009; 25(16):2078–2079.

21. Page AJ, Taylor B, Delaney AJ, Soares J, Seemann T, Keane JA, et al. SNP-sites: rapid efficient extraction of SNPs from multi-FASTA alignments. Microb genomics. 2016; 2(4).

22. Leigh JW, Bryant D. POPART: Full-feature software for haplotype network construction. Methods EcolEvol. 2015; 6(9):1110–1116.

23. GISAID. Clade and lineage nomenclature aids in genomic epidemiology studies of active hCoV-19 viruses [Internet]. 2020 [cited 2020 Jun 21]. Available from: https://www.gisaid.org/references/statements-clarifications/clade-and-lineage-nomenclature-aids-in-genomic-epidemiology-of-active-hCoV-19-viruses/

24. Korber B, Fischer WM, Gnanakaran S, Yoon H, Theiler J, Abfalterer W, et al. Tracking changes in SARS-CoV-2 Spike: evidence that D614G increases infectivity of the COVID-19 virus. Cell. 2020.

25. Zhang L, Jackson CB, Mou H, Ojha A, Rangarajan ES, Izard T, et al. The D614G mutation in the SARS-CoV-2 spike protein reduces S1 shedding and increases infectivity. bioRxiv. 2020; Forthcoming

26. Balaban M, Moshiri N, Mai U, Jia X, Mirarab S. TreeCluster: Clustering biological sequences using phylogenetic trees. PLoS One. 2019; 14(8):e0221068.

27. Kosakovsky Pond SL, Frost SDW. Not so different after all: A comparison of methods for detecting amino acid sites under selection. MolBiolEvol. 2005; 22(5):1208–1222.

28. Kosakovsky Pond SL, Frost SDW. Datamonkey: Rapid detection of selective pressure on individual sites of codon alignments. Bioinformatics. 2005; 21(10):2531–2533.

29. Cao H, Wang J, He L, Qi Y, Zhang JZ. DeepDDG: Predicting the Stability Change of Protein Point Mutations Using Neural Networks. J ChemInf Model. 2019; 59(4):1508–1514.

30. Gunalan, V., Mirazimi, A. & Tan, Y. AputativediacidicmotifintheSARS-CoVORF6proteininfluencesitssubcellularlocalizationandsuppressionofexpressionofco-transfectedexpressionconstructs. BMCResNotes. 2011; https://doi.org/10.1186/1756-0500-4-446

31. Hänel K, Willbold D. SARS-CoV accessory protein 7a directly interacts with human LFA-1. Biol Chem. 2007; 388(12).

32. Schaecher SR, Touchette E, Schriewer J, Buller RM, Pekosz A. Severe Acute Respiratory Syndrome Coronavirus Gene 7 Products Contribute to Virus-Induced Apoptosis. J Virol. 2007; 81(20):11054–11068.

33. Taylor JK, Coleman CM, Postel S, Sisk JM, Bernbaum JG, Venkataraman T, et al. Severe Acute Respiratory Syndrome Coronavirus ORF7a Inhibits Bone Marrow Stromal Antigen 2 Virion Tethering through a Novel Mechanism of Glycosylation Interference. J Virol. 2015; 89(23):11820–11833.

34. Zhang Y, Zhang J, Chen Y, Luo B, Yuan Y, Huang F, et al. The ORF8 Protein of SARS-CoV-2 Mediates Immune Evasion through Potently Downregulating MHC-I. bioRxiv. 2020; Forthcoming

35. Yoshimoto FK. The Proteins of Severe Acute Respiratory Syndrome Coronavirus-2 (SARS CoV-2 or n-COV19), the Cause of COVID-19. Protein J [Internet]. 2020;39(3):198–216.

36. Angeletti S, Benvenuto D, Bianchi M, Giovanetti M, Pascarella S, Ciccozzi M. COVID-2019: The role of the nsp2 and nsp3 in its pathogenesis. J Med Virol. 2020; 92(6):584–588.

