## Supplementary material for "In silico comparative genomics of SARS-CoV-2 to determine the source and diversity of the pathogen in Bangladesh": S1_SEQUENCES FROM BANGLADESH

**BANGLADESH ISOLATES**

| **GISAID ID** | **NAMING** | **COLLECTION DATE** | **ISOLATION LOCATION** |
| --- | --- | --- | --- |
| EPI_ISL_437912 | BS_1 | 4/18/2020 | Dhaka |
| EPI_ISL_445213 | BS_2 | 4/28/2020 | Dhaka |
| EPI_ISL_445214 | BS_3 | 4/28/2020 | Dhaka |
| EPI_ISL_445215 | BS_4 | 4/28/2020 | Dhaka |
| EPI_ISL_445216 | BS_5 | 4/28/2020 | Dhaka |
| EPI_ISL_445244 | BS_6 | 4/25/2020 | Dhaka |
| EPI_ISL_447904 | BS_7 | 5/11/2020 | Dhaka |
| EPI_ISL_450339 | BS_8 | 5/10/2020 | Chattogram |
| EPI_ISL_450340 | BS_9 | 5/13/2020 | Chattogram |
| EPI_ISL_450341 | BS_10 | 5/13/2020 | Chattogram |
| EPI_ISL_450342 | BS_11 | 5/8/2020 | Chattogram |
| EPI_ISL_450343 | BS_12 | 5/9/2020 | Chattogram |
| EPI_ISL_450344 | BS_13 | 5/3/2020 | Chattogram |
| EPI_ISL_450345 | BS_14 | 5/10/2020 | Chattogram |
| EPI_ISL_450839 | BS_15 | 5/6/2020 | Dhaka |
| EPI_ISL_450840 | BS_16 | 5/6/2020 | Dhaka |
| EPI_ISL_450841 | BS_17 | 5/13/2020 | Dhaka |
| EPI_ISL_450842 | BS_18 | 5/6/2020 | Dhaka |
| EPI_ISL_450843 | BS_19 | 5/6/2020 | Dhaka |
| EPI_ISL_455420 | BS_20 | 5/21/2020 | Dhaka |
| EPI_ISL_455458 | BS_21 | 5/21/2020 | Narayanganj |
| EPI_ISL_455459 | BS_22 | 5/21/2020 | Dhaka |
| EPI_ISL_458133 | BS_23 | 5/11/2020 | Dhaka |
| EPI_ISL_462090 | BS_24 | 5/23/2020 | Dhaka |
| EPI_ISL_462091 | BS_25 | 5/23/2020 | Dhaka |
| EPI_ISL_462092 | BS_26 | 5/23/2020 | Dhaka |
| EPI_ISL_462093 | BS_27 | 5/23/2020 | Dhaka |
| EPI_ISL_462094 | BS_28 | 5/23/2020 | Dhaka |
| EPI_ISL_462095 | BS_29 | 5/23/2020 | Dhaka |
| EPI_ISL_462096 | BS_30 | 5/23/2020 | Dhaka |
| EPI_ISL_462097 | BS_31 | 5/23/2020 | Dhaka |
| EPI_ISL_462098 | BS_32 | 5/21/2020 | Dhaka |
| EPI_ISL_464159 | BS_33 | 5/9/2020 | Narayanganj |
| EPI_ISL_464160 | BS_34 | 6/1/2020 | Dhaka |
| EPI_ISL_464161 | BS_35 | 6/1/2020 | Dhaka |
| EPI_ISL_464162 | BS_36 | 6/1/2020 | Dhaka |
| EPI_ISL_464163 | BS_37 | 5/31/2020 | Narayanganj |
| EPI_ISL_464164 | BS_38 | 5/26/2020 | Chattogram |
| EPI_ISL_464165 | BS_39 | 5/10/2020 | Narayanganj |
| EPI_ISL_464166 | BS_40 | 5/31/2020 | Narayanganj |
| EPI_ISL_465163 | BS_41 | 5/7/2020 | Narayanganj |
| EPI_ISL_465164 | BS_42 | 5/10/2020 | Narayanganj |
| EPI_ISL_466626 | BS_43 | 5/7/2020 | Narayanganj |
| EPI_ISL_466627 | BS_44 | 6/1/2020 | Narayanganj |
| EPI_ISL_466628 | BS_45 | 6/1/2020 | Narayanganj |
| EPI_ISL_466629 | BS_46 | 5/10/2020 | Narayanganj |
| EPI_ISL_466630 | BS_47 | 5/10/2020 | Narayanganj |
| EPI_ISL_466636 | BS_48 | 5/10/2020 | Narayanganj |
| EPI_ISL_466637 | BS_49 | 5/26/2020 | Chattogram |
| EPI_ISL_466638 | BS_50 | 5/26/2020 | Chattogram |
| EPI_ISL_466639 | BS_51 | 5/26/2020 | Chattogram |
| EPI_ISL_466644 | BS_52 | 6/1/2020 | Dhaka |
| EPI_ISL_466645 | BS_53 | 6/1/2020 | Dhaka |
| EPI_ISL_466649 | BS_54 | 6/1/2020 | Dhaka |
| EPI_ISL_466650 | BS_55 | 6/1/2020 | Dhaka |
| EPI_ISL_466686 | BS_56 | 5/9/2020 | Narayanganj |
| EPI_ISL_466687 | BS_57 | 5/26/2020 | Chattogram |
| EPI_ISL_466688 | BS_58 | 6/1/2020 | Dhaka |
| EPI_ISL_466689 | BS_59 | 6/1/2020 | Dhaka |
| EPI_ISL_466690 | BS_60 | 6/1/2020 | Dhaka |
| EPI_ISL_466691 | BS_61 | 6/1/2020 | Dhaka |
| EPI_ISL_466692 | BS_62 | 5/26/2020 | Chattogram |
| EPI_ISL_466693 | BS_63 | 6/1/2020 | Dhaka |
| EPI_ISL_466694 | BS_64 | 6/1/2020 | Dhaka |
