## Supplementary material for "In silico comparative genomics of SARS-CoV-2 to determine the source and diversity of the pathogen in Bangladesh": S2_SEQUENCES FROM SELECTED COUNTRIES

| **GISAID** | **ISOLATION DATE** | **COUNTRY** |
| --- | --- | --- |
| EPI_ISL_407894 | 1/28/2020 | Australia |
| EPI_ISL_408976 | 1/22/2020 | Australia |
| EPI_ISL_408977 | 1/25/2020 | Australia |
| EPI_ISL_410718 | 2/5/2020 | Australia |
| EPI_ISL_413594 | 2/28/2020 | Australia |
| EPI_ISL_414414 | 2/29/2020 | Australia |
| EPI_ISL_416411 | 1/25/2020 | Australia |
| EPI_ISL_416415 | 2/8/2020 | Australia |
| EPI_ISL_419733 | 1/31/2020 | Australia |
| EPI_ISL_419835 | 2/24/2020 | Australia |
| EPI_ISL_419945 | 3/21/2020 | Australia |
| EPI_ISL_426650 | 3/20/2020 | Australia |
| EPI_ISL_427643 | 2/28/2020 | Australia |
| EPI_ISL_430645 | 3/26/2020 | Australia |
| EPI_ISL_430714 | 4/10/2020 | Australia |
| EPI_ISL_436128 | 4/13/2020 | Australia |
| EPI_ISL_451120 | 3/31/2020 | Australia |
| EPI_ISL_451125 | 4/1/2020 | Australia |
| EPI_ISL_451531 | 3/28/2020 | Australia |
| EPI_ISL_451541 | 4/2/2020 | Australia |
| EPI_ISL_455057 | 5/1/2020 | Australia |
| EPI_ISL_455598 | 4/1/2020 | Australia |
| EPI_ISL_455603 | 1/30/2020 | Australia |
| EPI_ISL_456487 | 5/6/2020 | Australia |
| EPI_ISL_456615 | 5/19/2020 | Australia |
| EPI_ISL_456635 | 5/2/2020 | Australia |
| EPI_ISL_456646 | 5/26/2020 | Australia |
| EPI_ISL_412964 | 2/25/2020 | Brazil |
| EPI_ISL_413016 | 2/28/2020 | Brazil |
| EPI_ISL_414015 | 2/29/2020 | Brazil |
| EPI_ISL_414016 | 2/29/2020 | Brazil |
| EPI_ISL_415128 | 2/29/2020 | Brazil |
| EPI_ISL_417936 | 3/15/2020 | Brazil |
| EPI_ISL_427298 | 3/22/2020 | Brazil |
| EPI_ISL_429665 | 3/8/2020 | Brazil |
| EPI_ISL_429673 | 3/16/2020 | Brazil |
| EPI_ISL_450873 | 3/17/2020 | Brazil |
| EPI_ISL_456072 | 4/1/2020 | Brazil |
| EPI_ISL_456079 | 4/3/2020 | Brazil |
| EPI_ISL_456081 | 4/6/2020 | Brazil |
| EPI_ISL_456088 | 4/6/2020 | Brazil |
| EPI_ISL_456105 | 4/16/2020 | Brazil |
| EPI_ISL_413014 | 1/25/2020 | Canada |
| EPI_ISL_413015 | 1/23/2020 | Canada |
| EPI_ISL_415577 | 2/20/2020 | Canada |
| EPI_ISL_415578 | 2/27/2020 | Canada |
| EPI_ISL_415580 | 2/28/2020 | Canada |
| EPI_ISL_415581 | 3/1/2020 | Canada |
| EPI_ISL_418327 | 1/25/2020 | Canada |
| EPI_ISL_418344 | 2/29/2020 | Canada |
| EPI_ISL_418824 | 3/3/2020 | Canada |
| EPI_ISL_418834 | 3/11/2020 | Canada |
| EPI_ISL_418847 | 3/12/2020 | Canada |
| EPI_ISL_425177 | 1/23/2020 | Canada |
| EPI_ISL_429806 | 3/9/2020 | Canada |
| EPI_ISL_444501 | 4/6/2020 | Canada |
| EPI_ISL_444509 | 4/5/2020 | Canada |
| EPI_ISL_450640 | 4/8/2020 | Canada |
| EPI_ISL_450641 | 4/7/2020 | Canada |
| EPI_ISL_450644 | 4/8/2020 | Canada |
| EPI_ISL_450747 | 2/26/2020 | Canada |
| EPI_ISL_402120 | 1/1/2020 | China |
| EPI_ISL_402121 | 12/30/2019 | China |
| EPI_ISL_402124 | 12/30/2019 | China |
| EPI_ISL_406716 | 1/2/2020 | China |
| EPI_ISL_406717 | 1/2/2020 | China |
| EPI_ISL_408487 | 1/20/2020 | China |
| EPI_ISL_413875 | 2/10/2020 | China |
| EPI_ISL_413898 | 2/28/2020 | China |
| EPI_ISL_416326 | 1/30/2020 | China |
| EPI_ISL_416359 | 2/6/2020 | China |
| EPI_ISL_416403 | 2/2/2020 | China |
| EPI_ISL_421242 | 1/25/2020 | China |
| EPI_ISL_421259 | 1/27/2020 | China |
| EPI_ISL_428458 | 2/6/2020 | China |
| EPI_ISL_431118 | 3/13/2020 | China |
| EPI_ISL_434534 | 12/30/2019 | China |
| EPI_ISL_444969 | 4/16/2020 | China |
| EPI_ISL_450503 | 1/26/2020 | China |
| EPI_ISL_451312 | 1/26/2020 | China |
| EPI_ISL_451317 | 2/3/2020 | China |
| EPI_ISL_454942 | 3/2/2020 | China |
| EPI_ISL_454951 | 3/25/2020 | China |
| EPI_ISL_454957 | 3/19/2020 | China |
| EPI_ISL_454984 | 3/15/2020 | China |
| EPI_ISL_454987 | 3/2/2020 | China |
| EPI_ISL_455008 | 2/23/2020 | China |
| EPI_ISL_455011 | 2/11/2020 | China |
| EPI_ISL_455014 | 3/2/2020 | China |
| EPI_ISL_455363 | 3/2/2020 | China |
| EPI_ISL_455680 | 1/26/2020 | China |
| EPI_ISL_434680 | 4/8/2020 | Africa |
| EPI_ISL_408430 | 1/29/2020 | France |
| EPI_ISL_410486 | 2/8/2020 | France |
| EPI_ISL_410720 | 1/23/2020 | France |
| EPI_ISL_410984 | 1/29/2020 | France |
| EPI_ISL_411219 | 1/28/2020 | France |
| EPI_ISL_411220 | 1/28/2020 | France |
| EPI_ISL_414623 | 2/25/2020 | France |
| EPI_ISL_414624 | 2/26/2020 | France |
| EPI_ISL_414626 | 2/29/2020 | France |
| EPI_ISL_416746 | 3/3/2020 | France |
| EPI_ISL_418220 | 2/28/2020 | France |
| EPI_ISL_418225 | 3/8/2020 | France |
| EPI_ISL_418232 | 3/15/2020 | France |
| EPI_ISL_418422 | 3/17/2020 | France |
| EPI_ISL_420054 | 3/20/2020 | France |
| EPI_ISL_434629 | 4/7/2020 | France |
| EPI_ISL_443279 | 4/1/2020 | France |
| EPI_ISL_443291 | 4/2/2020 | France |
| EPI_ISL_443292 | 4/3/2020 | France |
| EPI_ISL_443307 | 4/9/2020 | France |
| EPI_ISL_406862 | 1/28/2020 | Germany |
| EPI_ISL_414505 | 2/27/2020 | Germany |
| EPI_ISL_414509 | 2/28/2020 | Germany |
| EPI_ISL_420900 | 3/12/2020 | Germany |
| EPI_ISL_425135 | 3/13/2020 | Germany |
| EPI_ISL_437204 | 3/5/2020 | Germany |
| EPI_ISL_437205 | 3/16/2020 | Germany |
| EPI_ISL_437207 | 4/9/2020 | Germany |
| EPI_ISL_437213 | 4/5/2020 | Germany |
| EPI_ISL_437226 | 4/10/2020 | Germany |
| EPI_ISL_437229 | 3/14/2020 | Germany |
| EPI_ISL_437284 | 4/2/2020 | Germany |
| EPI_ISL_437292 | 4/7/2020 | Germany |
| EPI_ISL_447610 | 2020-02 | Germany |
| EPI_ISL_447611 | 2020-02 | Germany |
| EPI_ISL_447613 | 2020-02 | Germany |
| EPI_ISL_450199 | 1/29/2020 | Germany |
| EPI_ISL_450200 | 1/28/2020 | Germany |
| EPI_ISL_450202 | 2020-01 | Germany |
| EPI_ISL_450209 | 1/30/2020 | Germany |
| EPI_ISL_413522 | 1/27/2020 | India |
| EPI_ISL_413523 | 1/31/2020 | India |
| EPI_ISL_420555 | 3/3/2020 | India |
| EPI_ISL_431103 | 3/16/2020 | India |
| EPI_ISL_435060 | 3/12/2020 | India |
| EPI_ISL_436451 | 4/13/2020 | India |
| EPI_ISL_447040 | 5/3/2020 | India |
| EPI_ISL_447559 | 3/31/2020 | India |
| EPI_ISL_447586 | 4/16/2020 | India |
| EPI_ISL_450325 | 3/17/2020 | India |
| EPI_ISL_452794 | 5/8/2020 | India |
| EPI_ISL_454560 | 4/6/2020 | India |
| EPI_ISL_455645 | 4/3/2020 | India |
| EPI_ISL_455653 | 4/21/2020 | India |
| EPI_ISL_455660 | 5/1/2020 | India |
| EPI_ISL_455767 | 5/7/2020 | India |
| EPI_ISL_458070 | 4/12/2020 | India |
| EPI_ISL_458095 | 5/24/2020 | India |
| EPI_ISL_424349 | 3/9/2020 | Iran |
| EPI_ISL_437512 | 3/26/2020 | Iran |
| EPI_ISL_442044 | 3/26/2020 | Iran |
| EPI_ISL_442523 | 3/9/2020 | Iran |
| EPI_ISL_445088 | 3/26/2020 | Iran |
| EPI_ISL_410545 | 1/29/2020 | Italy |
| EPI_ISL_410546 | 1/29/2020 | Italy |
| EPI_ISL_412974 | 1/29/2020 | Italy |
| EPI_ISL_413489 | 3/3/2020 | Italy |
| EPI_ISL_417921 | 3/1/2020 | Italy |
| EPI_ISL_417922 | 2/28/2020 | Italy |
| EPI_ISL_418256 | 3/14/2020 | Italy |
| EPI_ISL_420567 | 3/21/2020 | Italy |
| EPI_ISL_420569 | 3/23/2020 | Italy |
| EPI_ISL_420583 | 3/23/2020 | Italy |
| EPI_ISL_435151 | 4/8/2020 | Italy |
| EPI_ISL_435152 | 4/9/2020 | Italy |
| EPI_ISL_435153 | 4/9/2020 | Italy |
| EPI_ISL_435155 | 4/9/2020 | Italy |
| EPI_ISL_436727 | 4/27/2020 | Italy |
| EPI_ISL_436729 | 4/27/2020 | Italy |
| EPI_ISL_451298 | 2/12/2020 | Italy |
| EPI_ISL_451300 | 2/3/2020 | Italy |
| EPI_ISL_451301 | 2/3/2020 | Italy |
| EPI_ISL_451302 | 1/30/2020 | Italy |
| EPI_ISL_451306 | 2/21/2020 | Italy |
| EPI_ISL_451308 | 3/1/2020 | Italy |
| EPI_ISL_452186 | 3/28/2020 | Italy |
| EPI_ISL_452191 | 4/3/2020 | Italy |
| EPI_ISL_454733 | 3/1/2020 | Italy |
| EPI_ISL_457699 | 2/22/2020 | Italy |
| EPI_ISL_457728 | 3/4/2020 | Italy |
| EPI_ISL_457749 | 2/27/2020 | Italy |
| EPI_ISL_458084 | 4/3/2020 | Italy |
| EPI_ISL_458085 | 4/12/2020 | Italy |
| EPI_ISL_457839 | 2020-03 | Africa |
| EPI_ISL_457847 | 3/24/2020 | Africa |
| EPI_ISL_457848 | 3/24/2020 | Africa |
| EPI_ISL_457883 | 4/25/2020 | Africa |
| EPI_ISL_457884 | 4/25/2020 | Africa |
| EPI_ISL_457899 | 4/28/2020 | Africa |
| EPI_ISL_457910 | 4/8/2020 | Africa |
| EPI_ISL_412972 | 2/27/2020 | Mexico |
| EPI_ISL_424345 | 3/12/2020 | Mexico |
| EPI_ISL_424666 | 2/29/2020 | Mexico |
| EPI_ISL_424673 | 3/12/2020 | Mexico |
| EPI_ISL_424731 | 3/13/2020 | Mexico |
| EPI_ISL_426364 | 3/12/2020 | Mexico |
| EPI_ISL_452139 | 2/28/2020 | Mexico |
| EPI_ISL_455434 | 3/13/2020 | Mexico |
| EPI_ISL_458150 | 5/15/2020 | Africa |
| EPI_ISL_410301 | 1/13/2020 | Nepal |
| EPI_ISL_413550 | 2/27/2020 | Africa |
| EPI_ISL_455423 | 3/29/2020 | Africa |
| EPI_ISL_417444 | 3/4/2020 | Pakistan |
| EPI_ISL_419313 | 3/12/2020 | Pakistan |
| EPI_ISL_451958 | 3/16/2020 | Pakistan |
| EPI_ISL_427308 | 3/24/2020 | Russia |
| EPI_ISL_427310 | 3/29/2020 | Russia |
| EPI_ISL_427322 | 4/5/2020 | Russia |
| EPI_ISL_428893 | 3/23/2020 | Russia |
| EPI_ISL_428898 | 3/23/2020 | Russia |
| EPI_ISL_428899 | 3/29/2020 | Russia |
| EPI_ISL_428901 | 3/23/2020 | Russia |
| EPI_ISL_428910 | 3/31/2020 | Russia |
| EPI_ISL_428920 | 3/30/2020 | Russia |
| EPI_ISL_430071 | 4/8/2020 | Russia |
| EPI_ISL_430097 | 4/14/2020 | Russia |
| EPI_ISL_430108 | 4/15/2020 | Russia |
| EPI_ISL_450245 | 4/15/2020 | Russia |
| EPI_ISL_451968 | 3/18/2020 | Russia |
| EPI_ISL_416522 | 3/10/2020 | Saudi Arabia |
| EPI_ISL_437459 | 3/23/2020 | Saudi Arabia |
| EPI_ISL_437460 | 3/23/2020 | Saudi Arabia |
| EPI_ISL_437461 | 3/23/2020 | Saudi Arabia |
| EPI_ISL_437464 | 3/23/2020 | Saudi Arabia |
| EPI_ISL_437469 | 3/29/2020 | Saudi Arabia |
| EPI_ISL_437474 | 3/29/2020 | Saudi Arabia |
| EPI_ISL_437483 | 3/29/2020 | Saudi Arabia |
| EPI_ISL_437484 | 3/29/2020 | Saudi Arabia |
| EPI_ISL_437698 | 4/9/2020 | Saudi Arabia |
| EPI_ISL_437714 | 4/14/2020 | Saudi Arabia |
| EPI_ISL_437732 | 4/16/2020 | Saudi Arabia |
| EPI_ISL_437736 | 4/20/2020 | Saudi Arabia |
| EPI_ISL_437748 | 4/20/2020 | Saudi Arabia |
| EPI_ISL_437752 | 4/20/2020 | Saudi Arabia |
| EPI_ISL_437756 | 4/1/2020 | Saudi Arabia |
| EPI_ISL_437757 | 4/1/2020 | Saudi Arabia |
| EPI_ISL_437758 | 4/1/2020 | Saudi Arabia |
| EPI_ISL_437762 | 4/2/2020 | Saudi Arabia |
| EPI_ISL_418206 | 2/28/2020 | Africa |
| EPI_ISL_436686 | 3/27/2020 | Africa |
| EPI_ISL_455634 | 5/1/2020 | Africa |
| EPI_ISL_455635 | 5/5/2020 | Africa |
| EPI_ISL_455637 | 5/4/2020 | Africa |
| EPI_ISL_416483 | 2/26/2020 | Spain |
| EPI_ISL_419687 | 2/27/2020 | Spain |
| EPI_ISL_436294 | 3/24/2020 | Spain |
| EPI_ISL_436337 | 3/26/2020 | Spain |
| EPI_ISL_436361 | 2/29/2020 | Spain |
| EPI_ISL_436408 | 3/25/2020 | Spain |
| EPI_ISL_444972 | 4/5/2020 | Spain |
| EPI_ISL_447527 | 4/1/2020 | Spain |
| EPI_ISL_452365 | 4/10/2020 | Spain |
| EPI_ISL_452367 | 4/6/2020 | Spain |
| EPI_ISL_452378 | 3/12/2020 | Spain |
| EPI_ISL_452470 | 3/7/2020 | Spain |
| EPI_ISL_452615 | 4/1/2020 | Spain |
| EPI_ISL_452772 | 2/29/2020 | Spain |
| EPI_ISL_455351 | 2/29/2020 | Spain |
| EPI_ISL_428670 | 3/16/2020 | Sri Lanka |
| EPI_ISL_428671 | 3/10/2020 | Sri Lanka |
| EPI_ISL_428672 | 3/19/2020 | Sri Lanka |
| EPI_ISL_428673 | 3/31/2020 | Sri Lanka |
| EPI_ISL_450171 | 1/26/2020 | Sri Lanka |
| EPI_ISL_450172 | 2/8/2020 | Sri Lanka |
| EPI_ISL_430862 | 3/8/2020 | Sweden |
| EPI_ISL_454877 | 3/4/2020 | Sweden |
| EPI_ISL_454879 | 3/3/2020 | Sweden |
| EPI_ISL_454891 | 3/7/2020 | Sweden |
| EPI_ISL_455895 | 3/8/2020 | Sweden |
| EPI_ISL_427391 | 4/13/2020 | Turkey |
| EPI_ISL_428346 | 4/17/2020 | Turkey |
| EPI_ISL_428368 | 4/16/2020 | Turkey |
| EPI_ISL_429865 | 3/18/2020 | Turkey |
| EPI_ISL_435057 | 4/9/2020 | Turkey |
| EPI_ISL_437311 | 3/27/2020 | Turkey |
| EPI_ISL_437327 | 3/19/2020 | Turkey |
| EPI_ISL_437328 | 3/19/2020 | Turkey |
| EPI_ISL_455719 | 4/9/2020 | Turkey |
| EPI_ISL_457824 | 3/24/2020 | Turkey |
| EPI_ISL_451200 | 5/1/2020 | Africa |
| EPI_ISL_407071 | 1/29/2020 | UK |
| EPI_ISL_407073 | 1/29/2020 | UK |
| EPI_ISL_413221 | 2/28/2020 | UK |
| EPI_ISL_414005 | 2/25/2020 | UK |
| EPI_ISL_414006 | 2/28/2020 | UK |
| EPI_ISL_414012 | 2/27/2020 | UK |
| EPI_ISL_415133 | 2/29/2020 | UK |
| EPI_ISL_419440 | 3/17/2020 | UK |
| EPI_ISL_422023 | 3/30/2020 | UK |
| EPI_ISL_422037 | 3/27/2020 | UK |
| EPI_ISL_425463 | 3/7/2020 | UK |
| EPI_ISL_425856 | 3/18/2020 | UK |
| EPI_ISL_432244 | 4/4/2020 | UK |
| EPI_ISL_433151 | 4/16/2020 | UK |
| EPI_ISL_433532 | 3/30/2020 | UK |
| EPI_ISL_433556 | 4/1/2020 | UK |
| EPI_ISL_440338 | 3/20/2020 | UK |
| EPI_ISL_443781 | 3/30/2020 | UK |
| EPI_ISL_445467 | 4/3/2020 | UK |
| EPI_ISL_445541 | 4/5/2020 | UK |
| EPI_ISL_445625 | 4/7/2020 | UK |
| EPI_ISL_445726 | 4/10/2020 | UK |
| EPI_ISL_445734 | 4/13/2020 | UK |
| EPI_ISL_445846 | 4/10/2020 | UK |
| EPI_ISL_447942 | 5/1/2020 | UK |
| EPI_ISL_448066 | 5/5/2020 | UK |
| EPI_ISL_449004 | 4/19/2020 | UK |
| EPI_ISL_452867 | 5/11/2020 | UK |
| EPI_ISL_456669 | 5/5/2020 | UK |
| EPI_ISL_457560 | 5/11/2020 | UK |
| EPI_ISL_457612 | 4/14/2020 | UK |
| EPI_ISL_404253 | 1/21/2020 | USA |
| EPI_ISL_404895 | 1/19/2020 | USA |
| EPI_ISL_406034 | 1/23/2020 | USA |
| EPI_ISL_406036 | 1/22/2020 | USA |
| EPI_ISL_407214 | 1/25/2020 | USA |
| EPI_ISL_407215 | 1/25/2020 | USA |
| EPI_ISL_408670 | 1/31/2020 | USA |
| EPI_ISL_409067 | 1/29/2020 | USA |
| EPI_ISL_410044 | 1/27/2020 | USA |
| EPI_ISL_410045 | 1/28/2020 | USA |
| EPI_ISL_411954 | 2/6/2020 | USA |
| EPI_ISL_417111 | 3/5/2020 | USA |
| EPI_ISL_420585 | 3/18/2020 | USA |
| EPI_ISL_424900 | 3/7/2020 | USA |
| EPI_ISL_424907 | 3/6/2020 | USA |
| EPI_ISL_426461 | 3/27/2020 | USA |
| EPI_ISL_427528 | 3/12/2020 | USA |
| EPI_ISL_427538 | 3/13/2020 | USA |
| EPI_ISL_427618 | 3/16/2020 | USA |
| EPI_ISL_430190 | 3/16/2020 | USA |
| EPI_ISL_430960 | 4/1/2020 | USA |
| EPI_ISL_434133 | 3/28/2020 | USA |
| EPI_ISL_435625 | 3/6/2020 | USA |
| EPI_ISL_436468 | 3/26/2020 | USA |
| EPI_ISL_437384 | 4/16/2020 | USA |
| EPI_ISL_437389 | 4/13/2020 | USA |
| EPI_ISL_444060 | 4/27/2020 | USA |
| EPI_ISL_444656 | 3/30/2020 | USA |
| EPI_ISL_444740 | 4/2/2020 | USA |
| EPI_ISL_444744 | 4/2/2020 | USA |
| EPI_ISL_445177 | 3/31/2020 | USA |
| EPI_ISL_447214 | 3/18/2020 | USA |
| EPI_ISL_449824 | 4/7/2020 | USA |
| EPI_ISL_449826 | 4/7/2020 | USA |
| EPI_ISL_449860 | 4/8/2020 | USA |
| EPI_ISL_449890 | 4/12/2020 | USA |
| EPI_ISL_449996 | 4/16/2020 | USA |
| EPI_ISL_450086 | 3/12/2020 | USA |
| EPI_ISL_450128 | 3/19/2020 | USA |
| EPI_ISL_450175 | 4/4/2020 | USA |
| EPI_ISL_450847 | 4/9/2020 | USA |
| EPI_ISL_451234 | 5/4/2020 | USA |
| EPI_ISL_451235 | 5/4/2020 | USA |
| EPI_ISL_451241 | 5/1/2020 | USA |
| EPI_ISL_451247 | 5/4/2020 | USA |
| EPI_ISL_451275 | 5/7/2020 | USA |
| EPI_ISL_451277 | 5/6/2020 | USA |
| EPI_ISL_451284 | 5/6/2020 | USA |
| EPI_ISL_451295 | 5/6/2020 | USA |
| EPI_ISL_452108 | 3/10/2020 | USA |
| EPI_ISL_452121 | 3/23/2020 | USA |
| EPI_ISL_452798 | 4/3/2020 | USA |
| EPI_ISL_454687 | 5/12/2020 | USA |
| EPI_ISL_455577 | 5/20/2020 | USA |
